# Climate-driven increase in transmission of wildlife malaria parasite over the last quarter century

**DOI:** 10.1101/2025.03.25.644544

**Authors:** Angela Nicole Theodosopoulos, Fredrik Andreasson, Jane Jönsson, Johan Nilsson, Andreas Nord, Lars Råberg, Martin Stjernman, Ana Sofia Torres Lara, Jan-Åke Nilsson, Olof Hellgren

## Abstract

Climate warming is expected to influence the prevalence of vector-transmitted parasites. Understanding the extent to which this is ongoing, or has already occurred, requires empirical data from populations monitored over long periods of time, but these studies are sparse. Further, vector-disease research involving human health is often influenced by disease control efforts that supersede natural trends. By screening for malaria parasite infections in a wildlife population of blue tits (*Cyanistes caeruleus*) in Northern Europe, over a 26-year period, we tested whether observed prevalence and transmission changes were climate-driven and show that all three malaria parasite genera have increased significantly in their prevalence and transmission over time. The most common parasite in the study, *Haemoproteus majoris*, increased in prevalence from 47% (1996) to 92% (2021), and this is a direct consequence of warmer temperatures elevating transmission. Climate window analyses reveal that elevated temperatures between May 9^th^ and June 24^th^, a time period that overlaps with the host nestling period, are strongly positively correlated with *H. majoris* transmission in one-year-old birds. Warmer climate during this narrow timeframe has a demonstrable impact on parasite transmission, and this permeates into the overall prevalence in the host population. We now have empirical support that climate warming can drive a rapid rise in vector-transmitted parasites, and this has implications for other host-parasite systems. Given that we now know the exact time of year when climate warming is most influential on a common vector-transmitted parasite in this system, it is possible to investigate the evolutionary and environmental mechanisms that underly how these infections ultimately manifest. While more challenging to measure, similar implications of climate warming on human vector-disease systems might be occurring.

## Introduction

A rise in vector-borne diseases is a commonly expected effect that climate change can impose on ecosystems (Rocklöv & Dubrow, 2020; Semenza & Suk, 2018) but we lack long-term empirical evidence to support causation (Rocklöv & Dubrow, 2020, Chala & Hamde, 2021). For example, in 2010, an unprecedentedly warm year (Barriopedro et al., 2011), multiple Eurasian countries began seeing outbreaks of West Nile Virus (primarily transmitted by *Culex* mosquitoes) (Paz et al., 2013). While this outbreak was linked to elevated summer temperatures, the lack of extensive long-term data prevented pinpointing the direct causal effect of climate warming (Engler et al., 2013; Paz et al., 2013). The multifarious nature of climate change is at the helm of some vector population expansions (Cazelles et al., 2023; Duffy et al., 2024; Garamszegi, 2011; Turner et al., 2020). However, connecting climate-mediated vector trends with their associated host infection trends is challenging given the difficulty to disentangle effects of climate from other factors that can influence transmission (Barrero Guevara et al., 2024; Sadoine et al., 2024). Importantly, transmission is also affected by vector and host competence (Gervasi et al., 2015), habitat modifications such as urbanization (Wilke et al., 2019), and vector control efforts (Bhatt et al., 2015).

Since 1897, malaria parasites have played a pivotal role in research on vector-borne diseases. Mosquitoes were identified as a primary vehicle for their transmission to birds, and soon after, humans (Cox, 2010). Broadly, malaria parasites are defined as the parasitic alveolates that comprise the order Haemosporida. These parasites are found in a diversity of vertebrate hosts but the most common genera that infect birds are *Haemoproteus*, *Plasmodium*, and *Leucocytozoon*, each with their own set of vectors (biting midges, mosquitoes, and black flies, respectively; (Valkiunas, G., 2004)). As to humans, malaria, the disease that can result from haemosporidian parasite infections, can also be fatal to wildlife (LaPointe et al., 2012; Venkatesan, 2024). A key concern amongst epidemiologists and conservationists is whether climate change will drive a global rise in malaria infections (Cazelles et al., 2023; Mordecai, 2023; Sadoine et al., 2024), and this requires understanding the mechanisms that shape the prevalence and transmission of the parasite. We have already seen severe consequences from malaria on Hawaiian birds (LaPointe et al., 2012), and as malaria parasites often remain within hosts in the form of chronic infections (Valkiunas, G., 2004), a climate driven increase in their prevalence risks the accumulation of coinfections with more additive costs on host survival (Marzal et al., 2008).

There have been widespread control efforts targeting human malaria (Andrews et al., 1950; Bhatt et al., 2015; Duffy et al., 2024) making it difficult to pinpoint the extent to which a rise in ambient temperatures has already influenced parasite transmission, or how significant the effects of climatic changes are on the systems if left unchecked. As control efforts are lacking in even the most well-studied wildlife malaria systems (LaPointe et al., 2012) this enables their use for understanding how climate warming is shaping transmission. Importantly, few wildlife malaria investigations span more than a decade of data collection despite their potential significance for both human health and wildlife conservation at a global level (Bensch et al., 2007; Otero et al., 2019; Wilkinson et al., 2016). We are therefore severely limited in our understanding of how these parasites might have *already* increased in their prevalence given ongoing climate change. In Northern Hemisphere temperate zones, we can expect breeding seasons for many vectors and hosts to coincide during temporal windows governed by climatic conditions. As such, long-term studies targeting host breeding seasons in these regions, combining host-parasite infection data with climate and vector data, are especially suitable for understanding how vector-transmitted parasites are impacted by climate change. Yet, to our knowledge, there are no studies of this nature.

We sought to address the above knowledge gap by harnessing a long-term study of blue tits (*Cyanistes caeruleus*) in southern Sweden (Figure 1A). The blue tit is a small, typically non-migratory, passerine bird that is common in the Western Palearctic (Stenning, 2018). Blue tits nest in cavities, and as such, nest boxes can be deployed to study their breeding (Stenning, 2018). Since 1983 a population of blue tits has been studied within the Revinge area (Andreasson et al., 2023). Along with their breeding data, blood samples have been collected from birds every year since 1996. We used blood samples collected from three five-year periods, beginning in 1996 and ending in 2021, to identify malaria parasite infections. Based on data from the Swedish Meteorological and Hydrological Institute (*SMHI*, 2024), the temperature in the region has become significantly warmer since the mid-1990’s (Figure 1B). Climate change has been hypothesized to help facilitate a rise in many diseases resulting from vector-transmitted parasites, in part due to vectors being ectotherms that persist better in warmer conditions (Rocklöv & Dubrow, 2020). We therefore predicted that, over the course of the quarter century study period, we would see an overall increase in malaria parasite prevalence mediated by a climate-driven rise in transmission.

**Figure 1.**
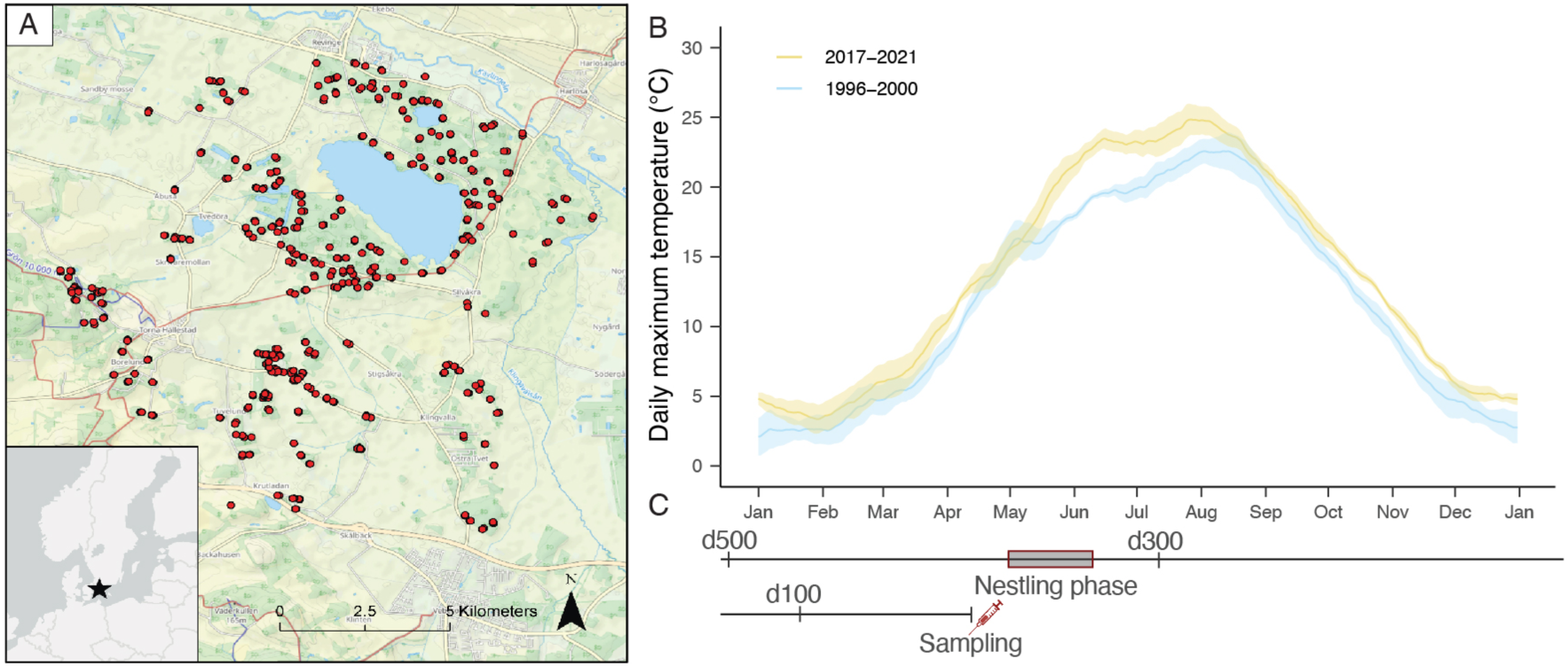
Between 1996 and 2021 blue tits were surveyed for malaria parasites at the Revinge field site in Southern Sweden at breeding sites indicated by red circles (A). 28 day rolling mean of the average maximum daily temperature for early and later years of the study. Ribbons represent ± one standard error (B). Birds were sampled during the nestling phase of their breeding season, and climate window analyses were allowed to extend 500 days prior to their sampling (d500, C).

## Materials and Methods

### Blue tit sampling

At the Revinge field site, blue tits (*Cyanistes caeruleus*) have been monitored since 1983 using a network of around 450 nest boxes (centered on 55.69°N and 13.46°E and approximately 30m in elevation) to study their breeding (Andreasson et al., 2023). Between the years 1996 and 2021 blood samples were collected from breeding adults between the months of April and July. All sampling was approved by the Malmö/Lund Animal Ethics Committee (permit nos. M 126-00; M 94-07; M 67-09; M 67-16; and 04705/2018). We conducted DNA extractions for *n =* 1966 samples, examined prevalence changes, and identified malaria parasite lineages using samples from 15 breeding seasons in three five-year time periods spanning 26 years: early (1996–2000, *n* = 472), middle (2007– 2011, *n* = 894), and later years (2017–2021, *n* = 600). Of all sampling events, 204 were from birds sampled more than once between field seasons. Throughout the duration of the study, hatching dates occurred between the 2^nd^ of May and the 5^th^ of June (Figure 2). Given that tit nestlings on average stay in the nest for 20 days (Nilsson & Svensson, 1996), the last fledglings will leave their nest on June 25^th^. Birds were aged based on plumage characteristics according to Svensson (1992) and sexed based on the presence or absence of a brood patch (Svensson, 1992).

**Figure 2.**
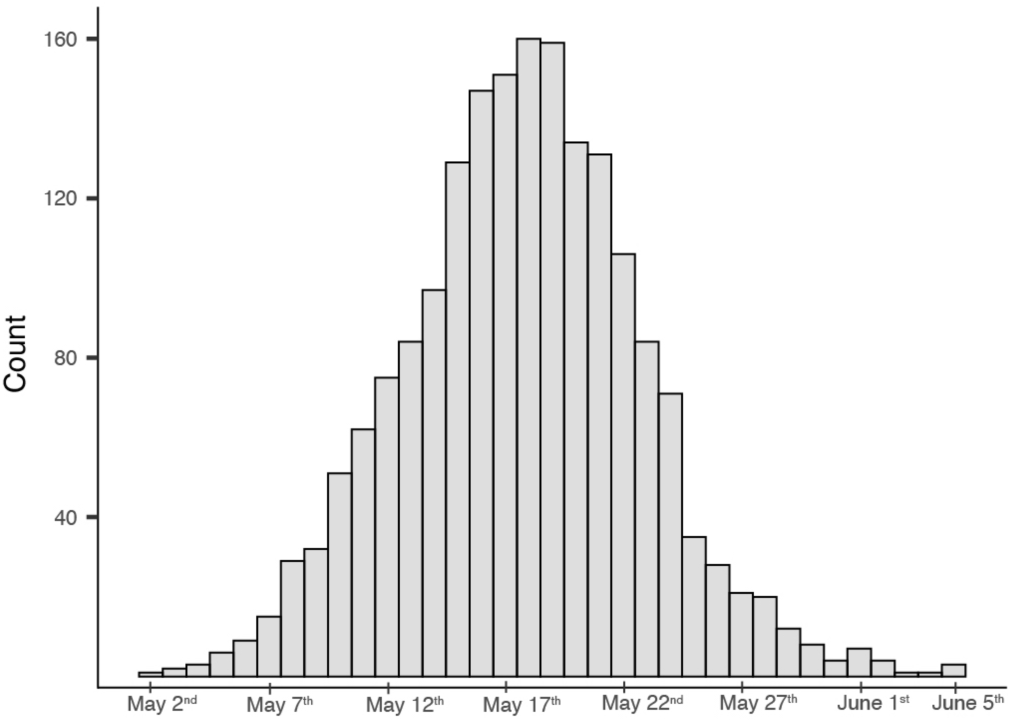
Histogram of hatching dates for all clutches hatched (n = 1882 clutches) spanning the three time blocks of the study. Hatching took place between May 2^nd^ and June 5^th^.

### Parasite screening

To simultaneously identify infections with *Haemoproteus*, *Plasmodium*, and *Leucocytozoon*, we used multiplex PCR methods (Ciloglu et al., 2019). All birds were screened twice to increase the detection probability of infections with low parasitemia (detail on lab work is available in extended methods). We included all samples from all breeding adults for assessing parasite prevalence because this best reflects the overall parasite load in the total population of blue tits at a given year, and in turn, the yearly parasite population. As such, including all annual samples enabled us to detect the overall accumulation or loss of parasites in the host population.

To examine prevalence, we used all *n* = 1966 samples from the blue tit population. However, birds often harbor low intensity chronic malaria parasite infections that can last multiple infection seasons (Knowles et al., 2010). Understanding temporal transmission trends therefore requires hosts to have only experienced a single season of parasite exposure upon sampling. We therefore used a subset of *n* = 1152 one-year-old birds (categorized as “Age 20”) to evaluate transmission. Thus, these individuals had only lived through one complete transmission year.

We used R (version 4.2.2) base functions to obtain the mean prevalence (proportion of individuals infected) for each year of the study (R Core Team, 2013). Additionally, we estimated prevalence, as an effect of year, by constructing generalized linear models with the “logit” link function and ran models separately for each malaria parasite genus. Models were also run separately for one-year-old birds to estimate transmission. To visualize models overlayed with yearly mean prevalence values we used the R packages *visreg* (Breheny & Burchett, 2017) and *ggplot2* (Wickham, 2011).

### Parasite typing

Variation in the mitochondrial *cytb* gene is commonly used to type genetic lineages of malaria parasites (Bensch et al., 2009). We typed genetic lineages using nested PCR methods, a commonly used practice, using previously described primers for simultaneous amplification of each of the three genera (Hellgren et al., 2004). PCRs were run separately to amplify (1) *Leucocytozoon*, and (2) *Haemoproteus* and *Plasmodium*. For all nested PCRs we included both negative controls, and positive controls for each of the three malaria parasite genera.

Nested PCRs were conducted using a subset of 657 samples for *Haemoproteus* and *Plasmodium* screening, and a subset of 83 samples for *Leucocytozoon* screening (see Supplementary Data). *Plasmodium* infections were challenging to type given that many were co-amplified with *Haemoproteus* which is usually present with a higher parasitemia in the blood (Waldenström et al., 2004). As such, we were only able to type 94 *Plasmodium* samples.

Following amplification of PCR products, we downloaded the resulting chromatogram files and uploaded them into Geneious software (Kearse et al., 2012). Finalized sequences were aligned to available typed lineage reads from the MalAvi database using the BLAST algorithm (Bensch et al., 2009; Johnson et al., 2008). To be assigned to a specific lineage the sequences needed to have a 100% match with previously described lineages from the MalAvi database. Some sequences could not be typed to lineage due to the presence of multiple mixed peaks observed in chromatograms, this was most common for *Leucocytozoon*. We tabulated all resulting lineages based on the individual that the lineage was sampled from, their age, and the year that they were sampled (Supplementary Data).

Molecular screening data and macroscopical examination of blood smears confirmed that low prevalence in early years is not due to sample degradation. As expected, PCR data nearly always presented almost equal or higher prevalence estimates than blood smears (Table 1), in contrast to what would be expected if DNA samples had degraded and failed to amplify (Stjernman, 2004).

**Table 1.**
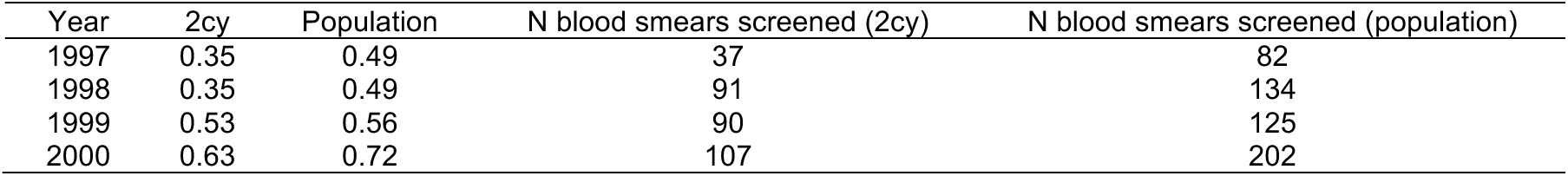
Blood smears were previously screened for *Haemoproteus* infections using samples from the same population to estimate prevalence (Stjernman, 2004). Importantly, these estimates show that PCR-methods are not underestimating prevalence during the early years which would have happened had the DNA samples degraded with time.

### Climate windows and path analyses

We conducted climate window analyses to detect potential climate signals for transmission of each of the three malaria genera (R package *climwin*: Bailey & Van De Pol, 2016; Van De Pol et al., 2016). The analyses compare the association between potentially important environmental variables in all possible time-windows within a selected timeframe and the response variable (i.e., parasite prevalence) using AICc (Akaike Information Criterion with small sample size correction; Burnham & Anderson, 2004). In our case we selected temperature as the environmental variable given that insect vectors, being ectotherms, rely heavily on their surrounding ambient temperature (Rocklöv & Dubrow, 2020).

We used daily mean, minimum, and maximum temperature recorded by a weather station (Swedish Meteorological and Hydrological Institute, station Lund 53430) approximately 20km from the study site as environmental predictors (i.e., climate signals) in a logistic regression model with yearly mean parasite prevalence of all one-year-old birds as the response variable. Each of the three parasite genera were modeled separately, using a binomial error distribution with the total number of sampled individuals each year added as a weighting factor. We allowed for both a linear and quadratic relationship between the environmental predictors and the response variable.

Birds were sampled during, or in conjunction with their breeding season in the spring (yearly median sampling date was between May 15^th^ and June 7^th^), and malaria parasites are not detectable until approximately 14 days after infection (Valkiunas, G., 2004). Therefore, we set the upper limit for the time-window to 14 days before the median date of the year with the latest median sampling date to reduce the possibility of detecting biologically irrelevant climate windows. Both biting midges (*Culicoides* spp.) and, to a lesser extent, black flies (Family: Simuliidae) are known to be present in nest-boxes during the nestling phase (Martínez-de La Puente et al., 2009). Therefore, we extended the timeframe of possible climate-windows back to January 1^st^ the preceding year, to include potential climate signals during the period before hatching and the nestling phase of sampled birds. Thus, we evaluated all climate-windows from May 24^th^, and extending back to January 1^st^ the preceding year. We also chose to exclude windows that were 10 days or shorter, as we considered such windows to be biologically implausible, and thereby increasing the risk for false positives if included.

Testing all possible time-windows within such a long timeframe increases the probability of finding a false positive, i.e., a signal that is not a true, biological signal but merely due to chance. Therefore, we ran 1,000 randomizations for each parasite-signal combination, where the randomizations remove any climate signal from the data and generates ΔAICc-values which can be compared to the ΔAICc-value of the best performing model. We used these comparisons to estimate the possibility that the climate window found is a false positive and not a true, biological signal. We only ran these randomizations for climate signals that were within two ΔAICc, as suggested in previous literature (Burnham & Anderson, 2004). We were interested in a stable estimate of the best performing climate signal and corresponding time-window. Therefore, we consistently used the median opening and closing dates of all windows that together make up 95% of the model weights, i.e., the 95% model confidence set, instead of the single best performing window (Table 2).

**Table 2.**
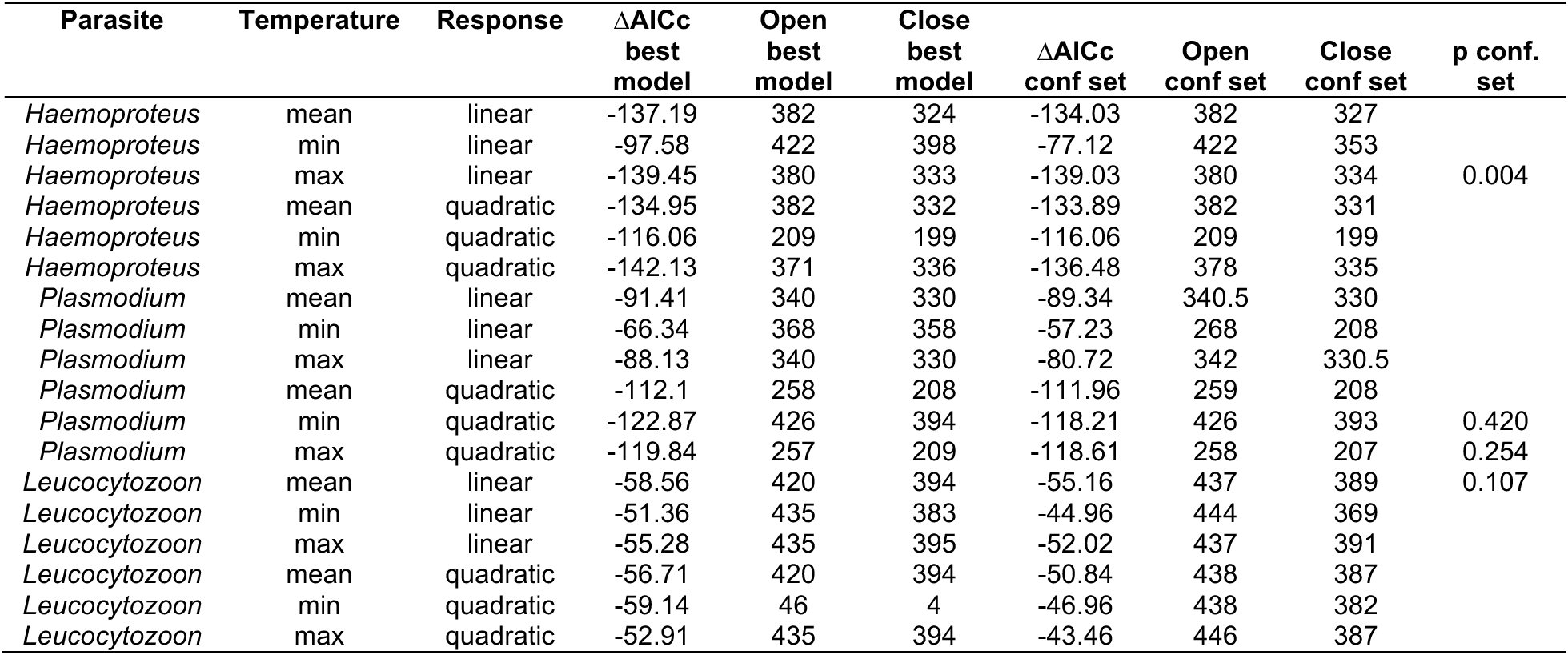
*Climwin* models tested for each malaria parasite (*Haemoproteus*, *Plasmodium*, and *Leucocytozoon*) using minimum, mean, and maximum daily temperatures and testing both a linear and quadratic relationship with response variables. ΔAICc with the best performing models are tabulated with their associated climate window opening and closing Julian dates. Following *climwin* documentation, we combined all windows that together make up 95% of the model weights (i.e., the 95% confidence set) with associated median opening and closing Julian dates, and probabilities (p conf set).

To tease apart the effect of temperature from any year effects unrelated to temperature we conducted a path analysis, using the R package *piecewiseSEM* (Lefcheck, 2016). The three paths estimated were (1) the effect of year on the temperature variable identified by *climwin*, (2) the effect of temperature on parasite prevalence and (3) the direct effect of year on parasite prevalence (unrelated to temperature). Parasite prevalence was modelled by logistic regression with climate window temperature and year as continuous predictors, and yearly mean parasite prevalence as the response variable, using a binomial error distribution with the total number of sampled individuals each year added as a weighting factor. The effect of year on climate window temperature was modelled by linear regression with a Gaussian error distribution. Standardization of coefficients was done using a latent-theoretical approach, as applied in *piecewiseSEM* (Grace et al., 2018). By uncoupling the effect of year from temperature, we were able to quantify how climate influenced parasite transmission. Since we were focused on the relationship between temperature and transmission, we only used data from one-year-old birds for all climate window and subsequent structural equation modeling analyses.

### Vector surveillance and identification

During the 2008 and 2009 field seasons vectors were surveilled from nest boxes at the Revinge field site (JÅN & MS, in preparation). Briefly, DNA was extracted from 30 of these individual vectors, all morphologically identified as biting midges, and molecular identification was done by amplifying the *cox1* gene using two different combinations of primers (Videvall et al., 2013). From the 30 samples, 17 had successful amplification of the mitochondrial *cox1* gene and were identified to species by aligning the resulting Sanger sequencing data to the NCBI database using the BLAST algorithm (Johnson et al., 2008).

## Results

### Lineages and prevalence changes

All *Haemoproteus* samples were of the same genetic lineage (PARUS1) belonging to the morpho-species *Haemoproteus majoris* (Nilsson et al., 2016). The majority of identified *Plasmodium* infections were either TURDUS1 (57%) or BT7 (42%), both of which belong to the same species, *Plasmodium circumflexum* (Valkiūnas et al., 2022). *Leucocytozoon* was the most diverse genus with at least eight different genetic lineages (not all infections could be identified to lineage from molecular methods due to multi-lineage coinfections; Supplementary Data Table 4).

We evaluated prevalence patterns spanning the study period using generalized linear models. *Haemoproteus* experienced a significant prevalence increase (p < 2.0 x 10^-16^) where prevalence rose from 47% (1996) to 92% (2021, Table 3). *Plasmodium* and *Leucocytozoon* showed more prevalence fluctuations during early and middle sampling years, but a consistently higher prevalence during later years (Table 3). As such, infections with *Plasmodium* and *Leucocytozoon* were significantly more common in later years (p = 4.7 x 10^-16^ and p = 2.3 x 10^-6^, respectively).

**Table 3.** Tabulation of *Haemoproteus* (HAEM), *Plasmodium* (PLAS), and *Leucocytozoon* (LEUCO) prevalence for one-year-old (2cy) birds and the total population for each year of sampling.

| Year | HAEM 2cy | PLAS 2cy | LEUCO 2cy | HAEM Population | PLAS Population | LEUCO Population |
| --- | --- | --- | --- | --- | --- | --- |
| 1996 | 0.41 | 0.93 | 0.76 | 0.47 | 0.95 | 0.76 |
| 1997 | 0.45 | 0.41 | 0.86 | 0.51 | 0.65 | 0.87 |
| 1998 | 0.49 | 0.27 | 0.59 | 0.58 | 0.38 | 0.59 |
| 1999 | 0.64 | 0.68 | 0.91 | 0.63 | 0.66 | 0.91 |
| 2000 | 0.66 | 0.56 | 0.76 | 0.71 | 0.60 | 0.78 |
| 2007 | 0.81 | 0.67 | 0.61 | 0.83 | 0.73 | 0.65 |
| 2008 | 0.85 | 0.36 | 0.91 | 0.87 | 0.46 | 0.85 |
| 2009 | 0.88 | 0.57 | 0.75 | 0.92 | 0.65 | 0.80 |
| 2010 | 0.70 | 0.66 | 0.87 | 0.77 | 0.68 | 0.88 |
| 2011 | 0.80 | 0.53 | 0.70 | 0.87 | 0.67 | 0.77 |
| 2017 | 0.95 | 0.86 | 0.80 | 0.96 | 0.90 | 0.80 |
| 2018 | 0.92 | 0.84 | 0.85 | 0.94 | 0.86 | 0.86 |
| 2019 | 0.96 | 0.79 | 0.85 | 0.96 | 0.86 | 0.83 |
| 2020 | 0.86 | 0.80 | 0.92 | 0.89 | 0.84 | 0.90 |
| 2021 | 0.86 | 0.81 | 0.96 | 0.92 | 0.89 | 0.91 |

### Transmission changes

For *Haemoproteus*, the temporal increase in transmission was equivalently significant for one-year-old birds as the prevalence increase for the broader population (p < 2.0 x 10^-16^, Figure 3A). *Plasmodium* transmission also increased significantly during the study period (p = 1.2 x 10^-12^, Figure 3B), as did *Leucocytozoon* (p = 6.9 x 10^-6^, Figure 3C).

**Figure 3.**
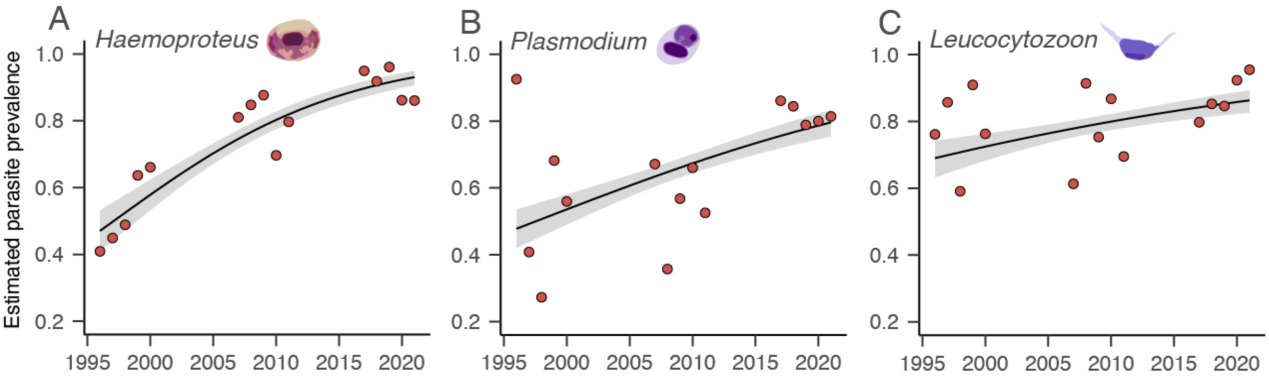
The annual predicted prevalence in relation to year for *Haemoproteus* (A), *Plasmodium* (B), and *Leucocytozoon* (C) in one-year-old birds from: logit(parasite genus) = *α* + *β*(Year) and with 95% confidence intervals shown. Points overlaying the model estimates and show the mean prevalence for each year.

### Relationship between transmission and temperature

*Haemoproteus* transmission had the highest supporting climate signal of all three genera (99.6% support, Figure 4A & B), based on the average daily maximum temperature in a window spanning May 9^th^ to June 24^th^ of the year before sampling. Notably, the climate window overlaps the nestling period for hosts in the study system almost perfectly (May 2^nd^ to June 25^th^), when one-year-old birds hatched the previous year (Figure 2) and were still residing in the nest boxes 20 days later (Figure 1C). Climate window temperature increased (± SE) with 0.17°C (± 0.04) for each year (t_13_ = 4.1, p = 1.2 x 10^-3^) and both temperature (z_12_ = 4.1, p < 1.0 x 10^-4^) and year (z_12_ = 3.8, p < 2.0 x 10^-4^) had a direct positive effect on parasite transmission (Figure 4C). The odds of getting infected increased with 39.0% for every °C increase in temperature (log odds ± SE: 0.329 ± 0.080, Figure 4D). The odds of getting infected also increased with 5.9% each year (log odds ± SE: 0.057 ± 0.015, i.e., independently of temperature). Interestingly, the direct effect of temperature was approximately 30% stronger compared to the direct effect of year (standardized coefficients of 0.32 vs. 0.24, Figure 4C & D). Thus, a higher maximum daily temperature during the nestling stage correlated with higher *Haemoproteus* transmission.

**Figure 4.**
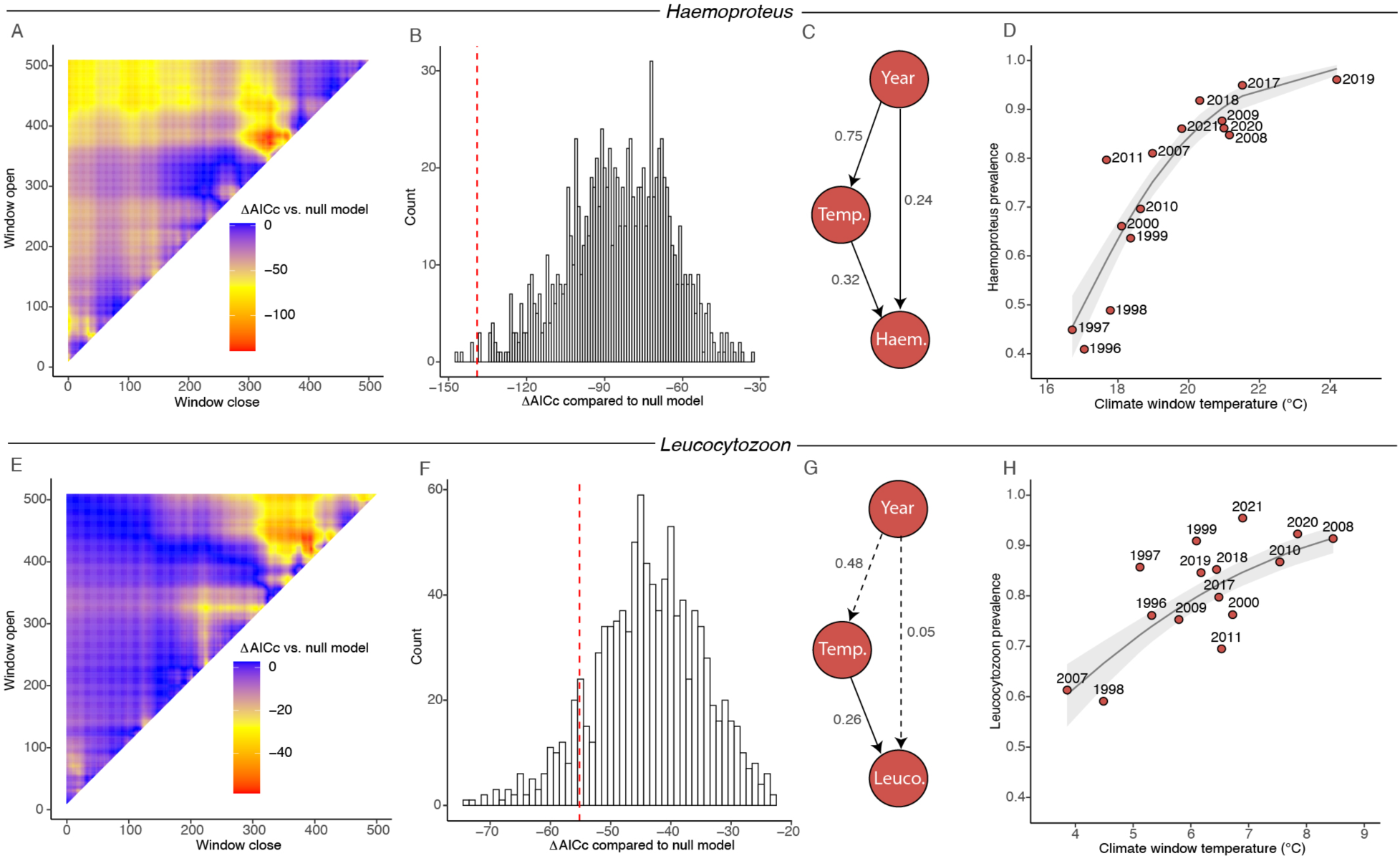
ΔAICc from *Haemoproteus climwin* analysis for the selected climate signal compared to the null model across all possible windows with the most likely window opening on May 9^th^ and closing June 24^th^ (A). ΔAICc of randomized data (histogram) compared to the obtained ΔAICc from the model shown as dashed red line (B). The structural equation model with standardized path coefficients (C). Yearly *Haemoproteus* prevalence in one-year-old birds as a function of climate window temperature (D). Dots are raw values, and the smoothed line indicates predicted probabilities together with their 95% confidence intervals. For E–H, the same outputs are shown for *Leucocytozoon* but with the most likely climate window opening on March 13^th^ and closing on April 30^th^. Dashed lines in g indicate non-significant (p > 0.05) effects.

For *Leucocytozoon* transmission, the best window had 89.3% support for being a true signal and was based on the average daily mean temperature with a window spanning March 13^th^ to April 30^th^ during the previous year (Figure 4E & F). Climate window temperature had a direct effect on parasite transmission (z_12_ = 6.0, p < 1.0 x 10^-4^) where the odds of getting infected increased with 48.1% for every °C increase in temperature (log odds ± SE: 0.393 ± 0.065, Figure 4G). During this window, the mean daily temperature tended to increase with year (0.07°C ± 0.03; t_13_ = 2.0, p = 0.067) but there was no direct effect of year on parasite transmission (z_12_ = 1.0, p = 0.33). *Leucocytozoon* had notably low prevalence in 1998 and 2007 (Figure 4H) and both these breeding seasons were preceded by relatively cold years, when transmission would have taken place. The two best supported climate windows for *Plasmodium* had low probabilities of being true signals (58 and 75%, respectively, Table 2.) and were therefore not analyzed further.

### The link to vectors

Climate window analyses indicated that *Haemoproteus* transmission occurs when blue tits are nestlings, therefore, *Haemoproteus*-transmitting vectors must be entering the nest boxes. Using molecular identification based on the *cox1* gene, we identified a subset of 17 individual midges to species. All three species found (*C. pictipennis*, *C. kibunensis* and *C. segnis*) in the nest boxes are presumed vectors for *H. majoris* (Chagas et al., 2024). Therefore, we can confirm that transmission is likely taking place when birds are nestlings.

Differences in biting midge abundance can vary annually, and this likely influences *Haemoproteus* prevalence in one-year-old birds the following year. We see evidence to support this in a notable drop in *Haemoproteus* prevalence between 2009 and 2010 (Figure 5A) corresponding to the reported lower biting midge abundance in 2009 compared to 2008 (JÅN & MS, in preparation; Figure 5B). Additionally, 2008 was a warmer year than 2009 (Figure 5C), consistent with a link between ambient temperature, the vector, transmission, and *Haemoproteus* prevalence.

**Figure 5.**
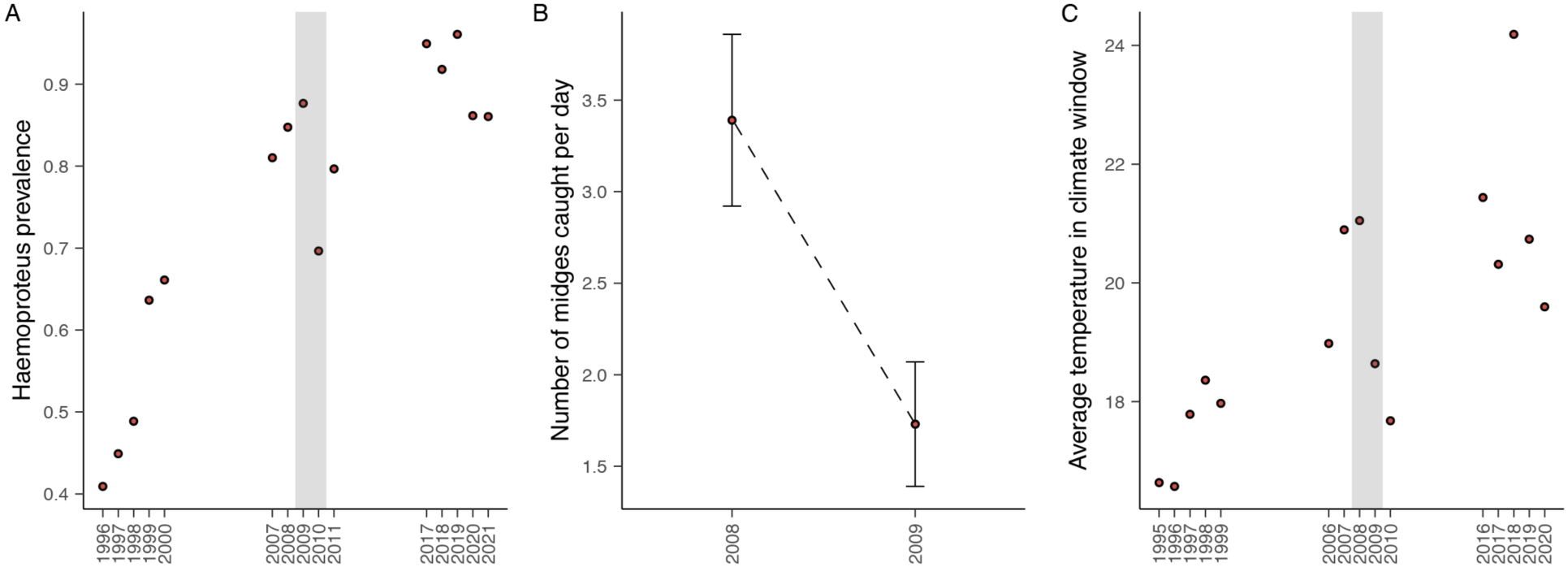
Average *Haemoproteus* prevalence for one-year-old birds spanning each year of sampling shows notably higher prevalence in the 2009 vs. 2010 breeding season (A). The number of *Haemoproteus*-transmitting biting midges recovered from nest boxes was notably higher in 2008 compared with 2009 (JÅN & MS, in preparation), supporting the prediction that more midges during the nestling period results in higher *Haemoproteus* prevalence the following year (B). Incidentally, the 2008 breeding season was warmer than in 2009, and this is in-line with our expectation that there are more *Haemoproteus-*harboring biting midges in warmer years than in cooler years (C).

## Discussion

Despite the ubiquity of parasites, our understanding of their infection trends rarely extends beyond a snapshot in time (Hayward et al., 2022) and this knowledge gap impairs our ability to predict how their prevalence and transmission will change as a result of ongoing global climate warming (Rizzoli et al., 2019; Rocklöv & Dubrow, 2020). Our study is the first to unambiguously show a causal effect of ambient temperature during a specific time window on the prevalence of a vector-transmitted parasite with indications that this is mediated by temperature effects leading to increased host-vector interactions. Unique to this study, a long-term natural history data set spanning over a quarter century, allowed us to follow the increase in parasite prevalence from moderate to high levels, through an increase in annual transmission. More one-year-old birds are getting infections in recent time, and this transmission increase permeates to a rise in prevalence for the total population of hosts.

For *Haemoproteus*, we now know that warmer temperatures during a short time window (between May 9^th^ and June 24^th^, when birds are nestlings) results in higher transmission and a subsequently higher prevalence, probably mediated by a temperature-dependent increased encounter rate with biting midges. Besides the general advantage of higher ambient temperatures for ectothermic vectors, knowledge of the specific climate window for parasite transmission enables us to also investigate specific mechanisms behind climate-driven transmission. For example, heat given off from hosts is known to attract vectors such as *Culex* mosquitoes (Reinhold et al., 2022) and birds have mechanisms to dissipate heat during warmer conditions which might, in turn, attract more vectors (Smit et al., 2016). Furthermore, nestling heat dissipation is costly (Smith et al. 2017) and will result in the production of more carbon dioxide, a trait that also attracts vectors (Castaño-Vázquez et al. 2020). Conducting field experiments that explore vector behavior as it relates to the rise in temperature and parasite prevalence can now be more targeted towards the relevant climate window. Furthermore, climate-driven increases in parasite transmission during a specific time window, may exert selection on breeding phenology and life-history traits in the host population. These factors need to be considered when predicting the resilience of populations in view of ongoing climate warming.

There are different climate window patterns for each of the three malaria parasite genera, which makes sense given their varied vectors and associated ecologies. Compared to *Haemoproteus*, the predicted climate window for *Leucocytozoon* lies earlier in the season (from March 13^th^ to April 30^th^, before eggs have hatched). The black flies (Family: Simuliidae) that transmit *Leucocytozoon* typically breed in flowing water (Adler & McCreadie, 2019) and the temperature of flowing streams at the Revinge field site might therefore play a role in determining when black flies emerge. An earlier vector emergence could mean more opportunities for *Leucocytozoon* transmission. *Leucocytozoon* is by far the most diverse of the malaria parasites that we sampled (at least eight different lineages). As such, there could be multiple relevant black fly species transmitting these parasites, thus making it harder to accurately predict how temperature is linked to vector ecology and *Leucocytozoon* transmission.

For avian *Plasmodium* parasites, mosquitoes of the genera *Culex*, *Culiseta*, *Aedes*, and *Anopheles* are all described vectors (LaPointe et al., 2012) but the specific species that transmit *Plasmodium circumflexum* (lineages TURDUS1 and BT7) are not currently known (Valkiūnas et al., 2022). We suspect that these vectors are actively transmitting infections during a relatively broad span of the blue tit annual cycle given our inability to identify any associated climate window.

In this study we have identified an effect of climate warming on the transmission and prevalence of malaria parasites in a population of blue tits. While this is only a single host-parasite system, similar impacts of climate change on vector-transmitted parasites might be occurring more broadly. For example, theoretical approaches have projected overall significant increases in vector-transmitted disease risk for humans with ongoing climate change (Colón-González et al., 2021; Martens et al., 1995; Ryan et al., 2021). Further, a key meta-analysis by Garamszegi (2011) showed a strong correlation between temperature anomalies and the increased incidence of avian malaria (Genus: *Plasmodium*). However, based off what we could infer from the literature cited by Garamszegi (2011), the studies included in this meta-analysis involved multiple host species and overwhelmingly lacked long-term data for each host population. Long-term data is necessary to reveal how parasite prevalence patterns in a host population are paralleled by corresponding climate changes. However, while more long-term studies are needed, it is important to realize that an overall rise in wildlife malaria prevalence, driven by climate warming, has already happened in our study population.

The effects of a rise in vector-mediated parasite transmission within a host population will depend on the extent to which the parasites are causing disease, and relatedly, the magnitude of host genetic variation for resistance and/or tolerance (Råberg et al., 2009). Importantly, variation in these traits will determine the rate of evolution for increased resistance/tolerance in the face of increased parasite prevalence (Roy & Kirchner, 2000). Understanding the factors that underly both parasite transmission, and host response to infection, will shed light on wildlife population resilience versus collapse given the ongoing rise of prevalence. As many vectors have highly similar ecologies, and the vectors responsible for transmitting wildlife parasites also might transmit disease-causing parasites to both livestock and humans (Rocklöv & Dubrow, 2020), further unravelling of their dynamics in relation to climate warming will be of utmost importance during the coming years.

## Acknowledgments

We thank the many people who assisted with data collection over the years of this study. Lab work was also made possible by the assistance and expertise provided by Tomas Johansson, Anna Sterngren, and Elsie Ye Xiong. Haemosporidian images were kindly provided by Tamara Emmenegger, and we thank Elana Engert for making the map of the Revinge field site. Funding for this work was provided by the Swedish Research Council through grants to OH (VR 2021-03663), to JÅN (VR 2021-05467), and to AN (VR 2020-04686). Support was also provided to OH through the Erik Philip-Sörensens Foundation (G2021-013), Kungliga Fysiografiska Sällskapet (2021/2068), and a fellowship provided by the Carl-Trygger Foundation (22:2338) that supported research by ANT.

## Extended Methods

### DNA extraction

From every bird sampled, 20-60µL of blood was preserved in 500µL of SET buffer (0.15M NaCl, 0.05M Tris, and 0.001M EDTA at pH ∼8.0). For DNA extraction, we combined ∼125µL of the SET buffer/blood mix with 2.5 µL of Thermo Scientific proteinase K (∼20 mg/ml), and 3.5µL of 20% SDS. Samples were then shaken and digested at 56°C overnight in a water bath. We then added 125µL of 4M NH_4_Ac to each sample, vortexed, and left samples at room temperature for 60 minutes where they were shaken and spun down every 15 minutes. Afterwards, we spun samples at 13000 rpms for 15 minutes to pellet all precipitate and remove the supernatant to a fresh tube. We then added 500µL of ice cold 95% EtOH to each sample, mixed thoroughly by shaking, then spun samples at 13000 rpms for 15 minutes. Following this procedure, we removed the supernatant and added an additional 250µL of ice cold 70% EtOH to the pelleted sample then immediately removed the liquid. Samples were dried overnight then dissolved in 20µL of 1xTE buffer. We then quantified samples using a nanodrop, and diluted samples to 25ng/µL for downstream PCR methods.

### Multiplex PCR

We used multiplex PCR methods to screen for the presence or absence of infection with each of the three malaria parasite genera (Ciloglu et al., 2019). First, we combined 5µL of Qiagen Multiplex Mastermix™, with 1.8µL of ddH_2_O and 2µL of 25ng/µL DNA template, with 0.2µL of each primer as described in previous literature (Ciloglu et al., 2019). We ran the reaction using the following thermocycler profile: a single cycle at 95°C for 15 minutes, 35 cycles of 94°C (30 seconds), 59°C (90 seconds), and 72°C (30 seconds), followed by a single cycle of 72°C ran for 10 minutes. We used gel electrophoresis to identify infections with each genus based on the expected length of DNA fragments where 2.5µL of the final PCR product mixed with 2µL of stopmix (0.01M EDTA, 15% Ficoll, 0.25% bromophenol, and 0.25% xylene cyanol FF) was ran in a 2% 96-well agarose gel combined with 15µL of Biotium GelRed solution. Gels were run at 80V for a maximum of 55 minutes. All multiplex PCRs were carried out in 96-well plates, with positive controls for each of the three genera, and negative controls. Infection scoring was based on both screens. Low intensity infections can have a “blinking on and off” effect with PCR screens (Hellgren et al., 2004). We therefore scored birds as infected if at least one screening revealed an infection. For samples that did not reveal clear results during their screening (i.e., bands appeared too faint to confidently score) they were scored as “NA.”

### Nested PCR

We first combined 1.0µL of 25ng/µL of DNA template with the following: 15.4µL of ddH_2_O, 1.5µL of MgCl_2_, 2.5µL of 10x reaction buffer, 2.5µL of 10x dNTP mix, 1µL of each 10 µM primer, and 0.1µL of Taq polymerase. Initial PCRs were run using the following thermocycler profile: a single cycle at 94°C for two minutes, 22 cycles at 94°C (30 seconds), 50°C (30 seconds), and 72°C (45 seconds), and ending with a single cycle at 72°C for 10 minutes. The second PCR was then conducted using 2µL of initial PCR product combined with the same concentrations of reagents as above (except we reduced the quantity of ddH_2_O to 14.4 µL). Additionally, we used the nested set of primers and ran the reaction for 35 cycles in the Thermocycler.

### *Cytb* sequencing

Amplified PCR products from nested PCRs were prepared for sequencing using an ABI 3100 Sanger Sequencer in the following two steps: (1) Precipitation of PCR products and (2) BigDye™ sequencing preparation. To initially precipitate PCR products we combined the 25µL of PCR product with 11µL of 8M NH_4_Ac and 37.5µL of 95% room temperature EtOH. Samples were then shaken and rested at room temperature for 15 minutes. We then pelleted samples by spinning at 3300rpms at 4°C for 30 minutes. After removing the supernatant, we added 50µL of ice cold 70% EtOH and immediately removed the supernatant followed by spinning the samples upside down at 3000rpms and at 4°C for one minute. We then dissolved the precipitated products in 25µL of ddH_2_O.

Sequencing preparation was conducted by combining 2µL of precipitated PCR product with the following: 5.0µL of ddH_2_O, 1.5µL of 5x Buffer, 0.5µL of the forward nested PCR primer (10µM), and 1µL of Applied Biosystems BigDye™ Terminator Ready Reaction Mix. We then ran the reaction using the following thermocycler profile: a single cycle at 96°C for one minute, followed by 25 cycles of 96°C (10 seconds), 50°C (5 seconds), and (60°C for four minutes). We then precipitated the final products by combining the 10µL of amplified product with 2.5µL of EDTA (125mM) and 35µL of 95% of room temperature EtOH, shaken, then let rest for 15 minutes. Samples were then pelleted by spinning at 3300rpms and at 4°C for 45 minutes. After removing the supernatant, we added 50µL of 70% ice-cold EtOH then immediately removed the supernatant. We then removed all remaining supernatant by spinning the samples upside-down at 3000rpms and at 4°C for one minute.

## Supplementary Data Legends

**Table 1.** Tabulations of *Haemoproteus*, *Plasmodium*, and *Leucocytozoon* lineages identified based on the parasite’s mitochondrial *cytb* gene with the individual ringnumber, year of sampling, and age for each host. Some hosts were sampled more than once.

**Table 2.** Tabulation of all *Haemoproteus* lineages identified based on the parasite’s mitochondrial *cytb* gene with the individual ringnumber, year of sampling, and age for each host. Below is the total number of all *Haemoproteus* lineages.

**Table 3.** Tabulation of all *Plasmodium* lineages identified based on the parasite’s mitochondrial *cytb* gene with the individual ringnumber, year of sampling, and age for each host. Below is the total number of all *Plasmodium* lineages. Samples listed as “P_Unknown” could not be identified to lineage.

**Table 4.** Tabulation of all *Leucocytozoon* lineages identified based on the parasite’s mitochondrial *cytb* gene with the individual ringnumber, year of sampling, and age for each host. Below is the total number of all *Leucocytozoon* lineages. Samples listed as “Coinfection” could not be identified to lineage due to the presence of mixed peaks in chromatograms.

